# Clustering enzymes using *E.coli* inner cell membrane as scaffold in metabolic pathway

**DOI:** 10.1101/230425

**Authors:** You Wang, Yuqi Wu, Yang Suo, Huaqing Guo, Yineng Yu, Ruonan Yin, Rui Xi, Jiajie Wu, Nan Hua, Yuehan Zhang, Shaobo Zhang, Zhenming Jin, Yushu Wang, Lin He, Gang Ma

**Author notes:** To whom correspondence should be addressed. Dr. Ma Gang.

## Abstract

Clustering enzymes in the same metabolism pathway is a natural strategy to enhance the productivity. Several systems have been designed to artificially cluster desired enzymes in the cell, such as synthetic protein scaffold and nucleic acid scaffold. However, these scaffolds require complicated construction process and have limited slots for target enzymes. Following this direction, we designed a scaffold system based on natural cell membrane. Target enzymes (FabZ, FabG, FabI and TesA’ in fatty acid synthesis II pathway) are anchored on the *E.coli* inner membrane, showing the enhanced metabolism flux without the requirement of the further artificial interactions to force the clustering. Furthermore, anchoring the enzymes on the membrane enhances the products exportation, which further increases the productivity. Together, the proposed system has potential applications in producing valuable biomaterials.

## Background

In cells, many enzymes uptake energy and produce life essential materials. Some of them in the relevant metabolism flux naturally cluster as multienzyme “sequential” or “cascade” reactions such as glycolysis and Krebs cycle (*1, 2*). The existence of these natural “flow lines” indicates that clustering relevant enzymes could improve the efficiency of the metabolism flux and thus saves the energy in the cell.

To mimic the natural multienzyme complex, several approaches have been developed to organize functional related enzymes. These include designing artificial protein scaffold to generate desired metabolism flux (3). By using well-characterized and widespread protein-protein interaction domains from metazoan signaling proteins (SH3, PDZ and GBD binding domain), the authors produce a modular genetically encoded scaffold system: the localization of enzymes is predefined and programmable. Additionally, the authors show a 77-fold higher level of the products by using this system, demonstrating attractive applications of artificial scaffolds (3, 4).

Protein scaffold and other scaffold systems have predefined artificial scaffolds which generally are limited by the scaffold length or the number of modules *(5*, 6). Compared to the volume of cytoplasm, the cell membrane has a much smaller space, restricting membrane proteins in a certain region for different functions. The natural property of cell membrane make it become a potential scaffold to achieve clustering desired enzymes and enhance metabolism flux.

To verify this idea, we choose to anchor the key enzymes of *E.coli* fatty acid metabolic pathway into the inner cell membrane. *E.coli* fatty acid synthesis pathway contains nine enzymes: FabA, FabB, FabD, FabF, FabG FabH, FabI, FabZ, and ACP. One equivalent of acetyl-CoA and 6–8 equivalents of malonyl-CoA are converted into C14–C18 acyl-ACP species in this pathway (Fig. S1). Additionally, the Periplasmic thioesterase, TesA, is capable of releasing free fatty acids from acyl-ACP species (7). Previous studies suggested that fabG, fabI, fabZ and TesA control the rate-limiting steps in fatty acid over-production in *E.coli (7–9*). Based on these studies, we choose to fuse fabG, fabI, fabZ and TesA to Lgt, a well-studied *E.coli* inner membrane protein (*10*). By anchoring these enzymes, we observe increased final products yield and dramatically enhanced products exportation. Collectively, our results provide novel insight into the potential application of cell membrane as a scaffold for important metabolism pathway to produce valuable bioproducts.

## Materials and Methods

### Strains and constructs

*E. coli* DH5α was used for cloning and *E. coli* BL21 (DE3) were used for protein expression. The vectors for the constructs included pETDuet1, pACYCDuet1, pRSF-Duet1, and pBAD18.

The arabinose operon from pBAD18 was amplified and cloned twice into pETDuet1, pACYCDuet! or pRSF-Duet1, resulting in pET-Ara, pACYC-Ara or pRSF-Ara. Each of this vector contains two copies of arabinose operons. For the verification of membrane localization, the DNA fragment containing N-terminal DsbA signal sequence, followed by β-lactamase, Lgt and GFP was cloned into pET-Ara, under the control of arabinose promoter (Fig. 1A). For the verification of artificial clustering, the DNA fragment containing DsbA signal sequence, followed by split GFP, the cytoplasm protein interaction domain, Lgt and C-terminal periplasm protein interaction domain was cloned into arabinose operon (Fig. 2A and 2B). The DNA fragments of Group 1, Group2 and Group 3 proteins were cloned into pET-Ara, pACYC-Ara and pRSF-Ara, respectively. Control group genes were cloned into pET-Ara.

**Figure 1.**
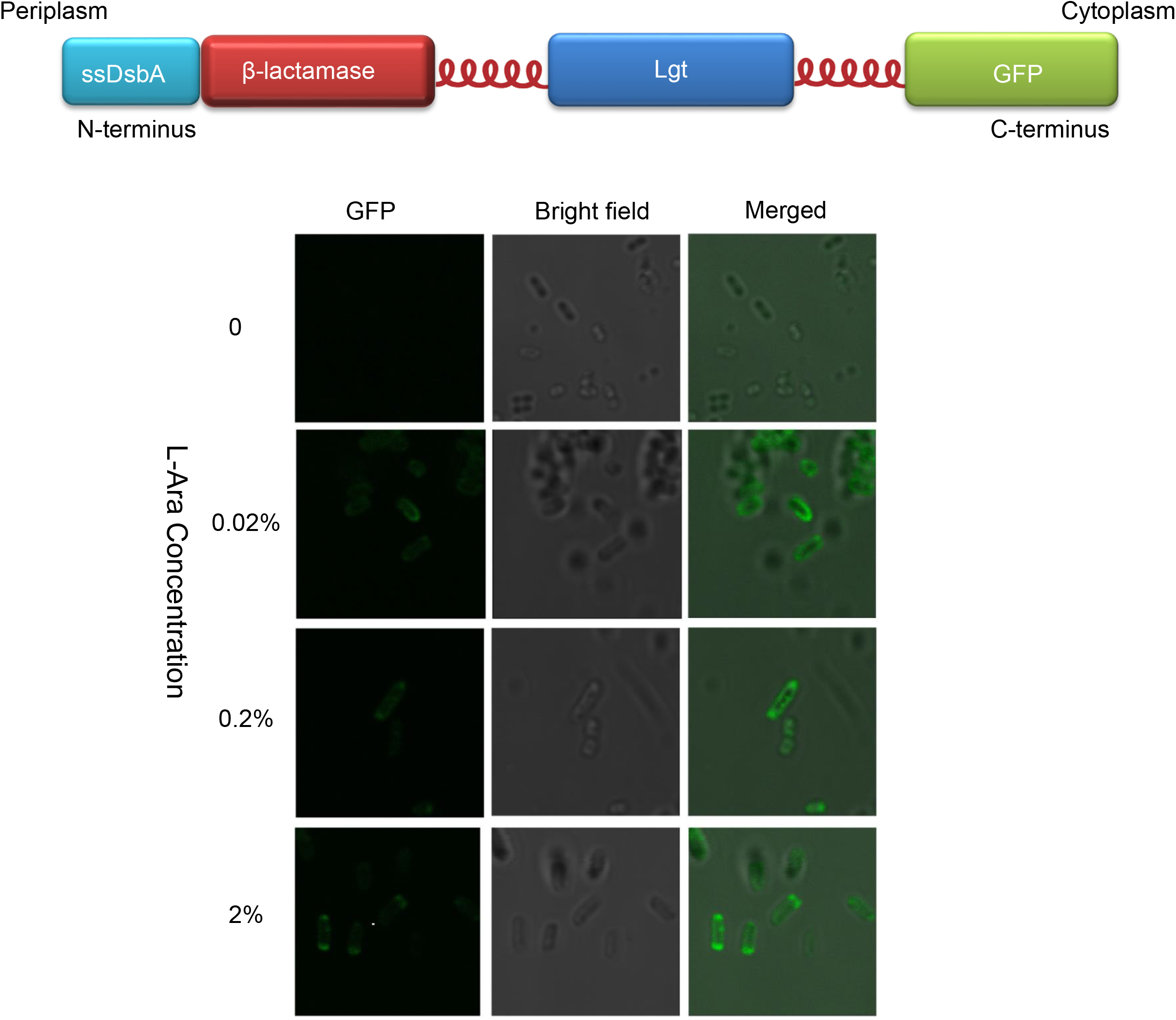
Design and verification of membrane localization of engineered Lgt. A) Schematic representation of engineered membrane protein. Lgt is used as a scaffold to carry functional groups (β-lactamase and GFP as examples) to the membrane. B) Membrane localization is verified by confocal microscopy.

**Figure 2.**
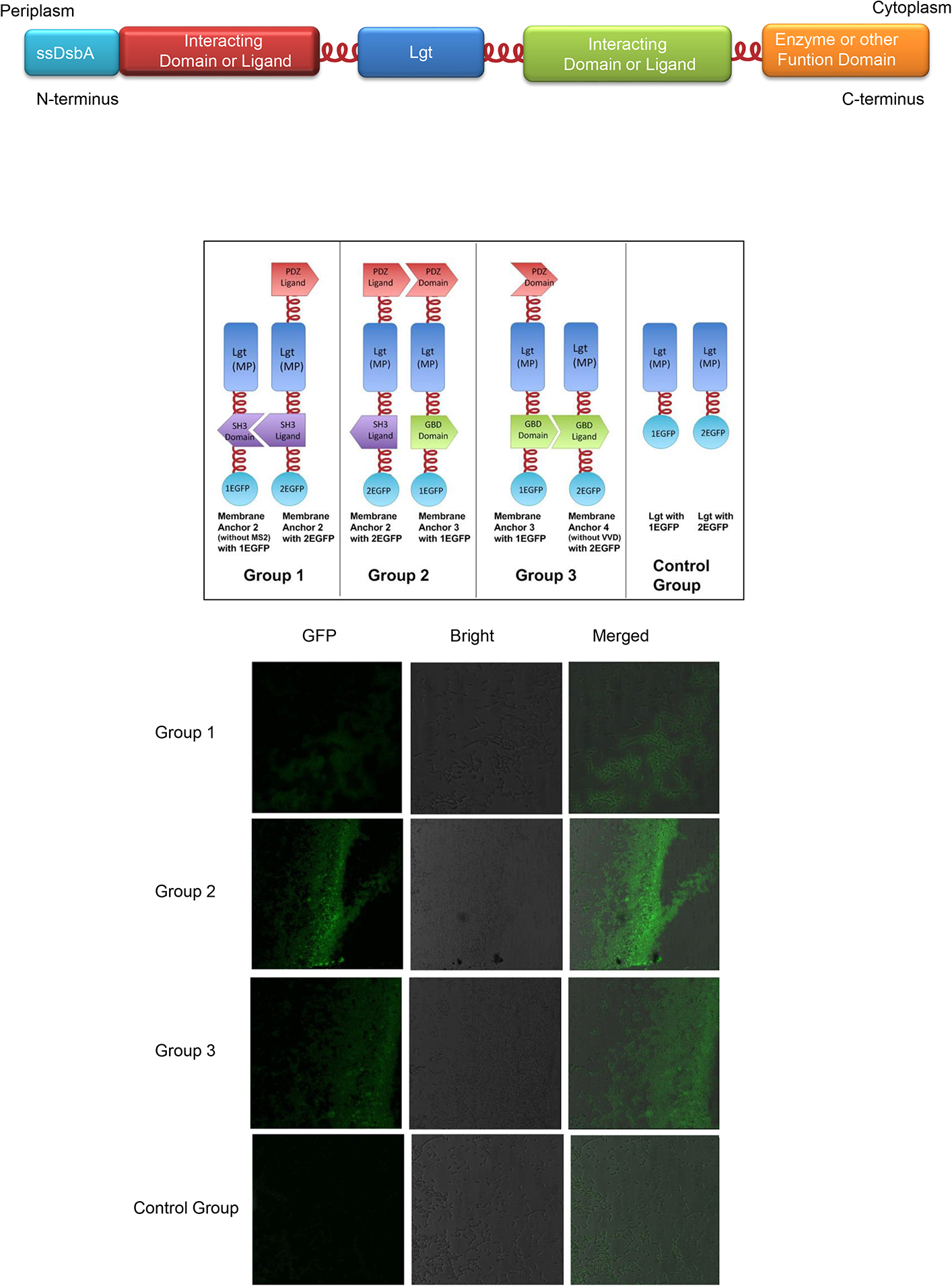
Design and verification of artificial membrane clustering. A) Schematic representation of engineered membrane protein. Protein interaction domains are fused to the ends of Lgt. B) Schematic representation of four groups of engineered membrane protein. Group1, 2 and 3 use SH3, PDZ and GBD interactions, respectively. Control group has no interaction domains. Split GFP is fused the N terminal of the membrane protein to verify the protein interactions. C) The protein interactions in different groups are verified by confocal microscopy. The detected GFP fluorescence indicates the interactions between designed proteins.

Four groups of engineered fatty acid related enzymes were used to verify our design (Fig. 4A). FabI, FabZ, FabG and TesA’ were fused to N terminal of each protein. Group1 contains cytoplasm and periplasm protein interaction domains to cluster engineered proteins. The enzymes in Group2 were directly fused to Lgt. The enzymes in Group3 were directly fused to protein interaction domains and expressed in the cytoplasm. The control group is the cytoplasm expressed enzymes. In each group, FabI and FabZ proteins or fusion proteins were cloned in two arabinose operons in pET-Ara, respectively. FabG and TesA’ proteins or fusion proteins were cloned in two arabinose operons in pRSF-Ara, respectively.

### Cell culture conditions for fatty acid biosynthesis

Cells carrying different constructs were incubated in 5 mL of LB medium supplemented with antibiotics and cultured overnight at 37 °C. Three percent (v/v) overnight cell culture were added to a 250 mL flask containing 50 mL of LB medium supplemented with 15 g·L^−1^ glucose and antibiotics and then cultivated at 37 °C under 150 rpm. The cultures were induced by addition of 0.2% arabinose at OD600 = 0.6, and samples were collected 20 h post-induction for fatty acid analysis.

### Free fatty acid extraction and analysis

Twenty-milliliter samples of cell culture (three replicates for each sample) were centrifuged at 8000□rpm for 10□min to separate cell-associated fatty acids from extracellular fatty acids. Fatty acid extraction was carried out as described earlier (*11*). The fatty acids extracted from the supernatant were analyzed by GC/MS using a 5975□C Series MSD and Agilent 6850 equipped with an HP-5□MS column (3□m□x□0.3□mm, film thickness of 0.25□mm). Helium was used as a carrier gas. The temperatures of the injector and detector were 250□°C and 280□°C respectively. The GC elution conditions were as follows: 100□°C as the starting temperature (for 5 min), 15 Jmin ramp to 250□°C, and 5 Jmin holding at 250□°C. All samples were spiked with pentadecanoic fatty acid (C15) as an internal standard. We repeated the growth of the cell and the analysis of the fatty acid products for three times.

## Results

### Localizing target enzymes to *E.coli* inner membrane

To utilize membrane as a scaffold to cluster target enzymes, we choose an *E.coli* membrane protein, Lgt as a membrane anchor. Target enzymes can be fused to Lgt N terminal as a periplasm enzyme, or Lgt C terminal as a cytoplasm enzyme. To verify this construct, we fused β-lactamase to the N terminal of Lgt and GFP to the C terminal (Fig.1A) (*12*). The expression plasmid containing this construct was transferred to BL21 (DE3) and the expression was induced by L-arabinose (Fig.1B). The confocal microscopic image of the cell shows that the fusion protein was correctly localized on the cell membrane. To verify whether the periplasm part of the fusion protein was properly presented, we tested cell viability in the presence of ampicillin and different concentration of L-arabinose (Fig. S2). The results show that the cell was alive only when the expression of the fusion protein was induced by L-arabinose. Together, these results show that the whole fusion protein had correct functions as what we designed.

### Clustering target enzymes on the membrane

Cell membrane has much smaller space compared to the cytoplasm volume, making us choose the *E.coli* inner cell membrane as a scaffold. However, it is unclear whether proteins anchored on the membrane can be clustered, showing similar property as other artificial scaffold to improve reaction flux. To answer this question, we first combined our membrane fusion protein and the protein scaffold (Fig. 2A and 2B) (*3*). Three groups of interaction proteins were fused to the N-terminal of the C-terminal of Lgt to create desired protein complexes on the membrane (Fig. 2B). To verify these interactions, we utilized the fluorescence complementation assay (*13*). The split GFP parts were fused to Lgt as negative control, or fused to Lgt interaction groups to test protein interactions (Fig. 2B). As shown in figure 2C, the detected GFP signal indicates that all three groups of Lgt fusion proteins formed desired complexes on the membrane. These results show that we construct a series of membrane protein complexes which can be used as the control to evaluate the property of the cell membrane as a scaffold to cluster target proteins.

### Clustering fatty acid synthesis enzymes on the membrane

To evaluate our design, we choose to apply the *E.coli* fatty acid synthesis pathway to our system. The key enzymes in this pathway, FabI, FabZ and FabG, and TesA were fused to the C-terminal of Lgt with (Fig. 3A) or without (Fig. 3B) the interaction protein groups. The consequence constructs have enzymes on the membrane and in the cytoplasm side. The first group of fusion proteins (Fig. 3A) are forced to be clustered by protein interactions. The proteins in the second group (Fig. 3B) are relative independent on the membrane. The cells expressing group 1 or group 2 proteins were cultured and the total fatty acids were extracted and measured (Fig. 3C). Compared to just overexpressing the enzymes in the cytoplasm, anchoring enzymes on the membrane (both group 1 and group 2) produced more fatty acid products. Comparing two membrane groups, the independent membrane enzymes produced slightly products than forced clustered enzymes. Our results support the idea that localizing on the membrane has similar clustering effects compared to the artificial protein scaffold, resulting in increased final products yield.

**Figure 3.**
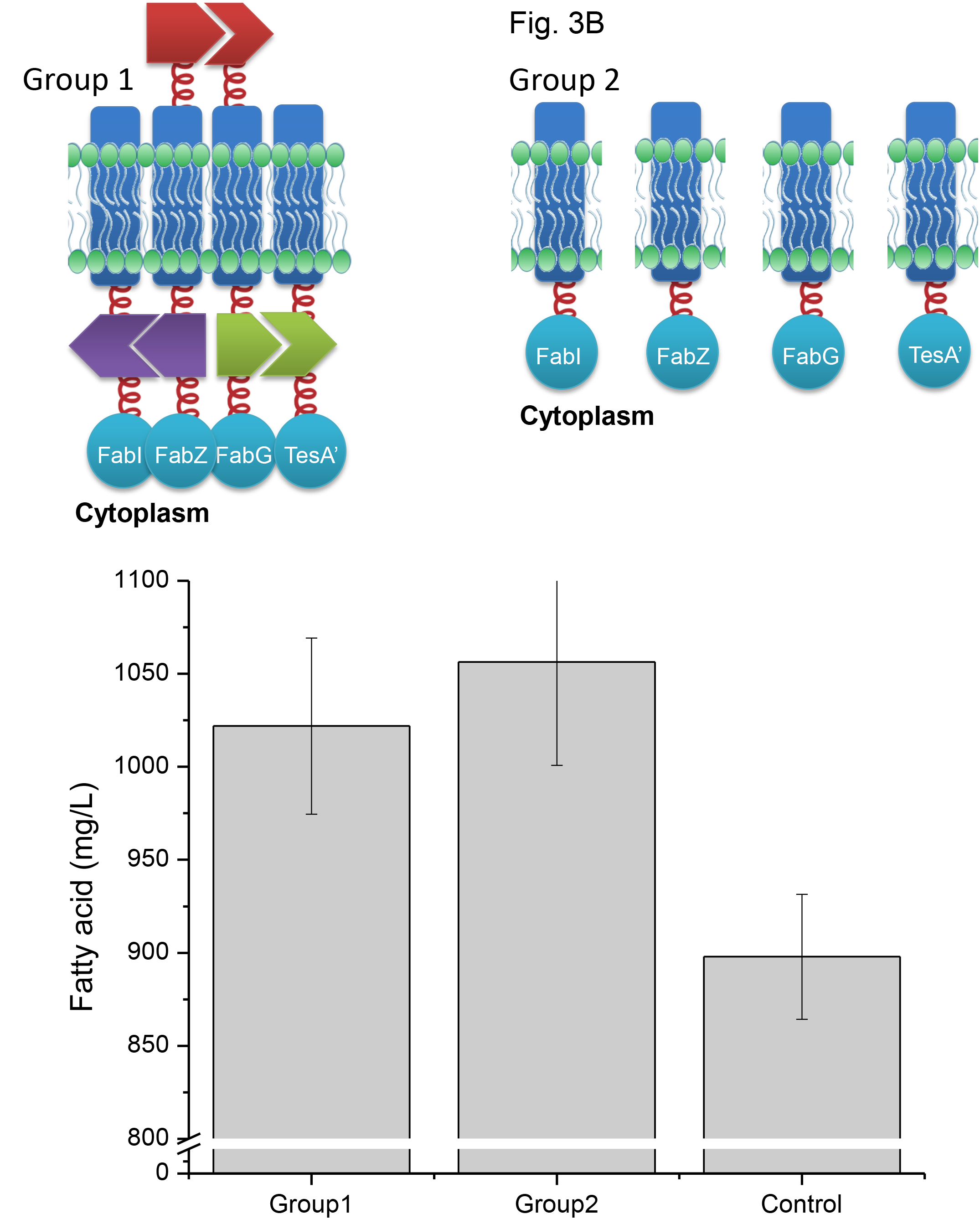
Clustering fatty acid metabolism enzymes on the membrane enhances the products yield. A) Schematic representation of two strategies to cluster enzymes. Group1 uses protein interaction domains to cluster target enzymes. Group2 has only Lgt domain to localize target enzymes on the membrane. B) Total fatty acid extracted from the cell expressing each groups of engineered enzymes. The control group is only expressing the four enzymes in the cytoplasm.

### Clustering fatty acid synthesis enzymes on the membrane accelerates products exportation

When we analyzed the fatty acid products results, we noticed that by anchoring enzymes on the membrane dramatically change the ratio between the products in the cell and in the medium. To further investigate this phenomenon, we further constructed the clustered enzymes in the cytoplasm (group 3) and compared the fatty acid products yield between four groups (Fig. 4A). We found that the membrane groups (group1 and group 2) exported more fatty acid products than the cytoplasm groups (group 3 and control) (Fig. 4B). Comparing clustering enzymes on the membrane (group 1 and group 2) with in the cytoplasm (group 3), we found that anchoring enzymes on the membrane produced more total fatty acid products (Fig. 4C). Clustering enzymes in the cytoplasm (group 3) had a higher yield than control (Fig. 4C), which is consistent with protein scaffold studies (*3*). Together, our results show that anchoring enzymes on the membrane can also enhance the products exportation, may resulting in the further higher yield than just clustering enzymes in the cytoplasm.

**Figure 4.**
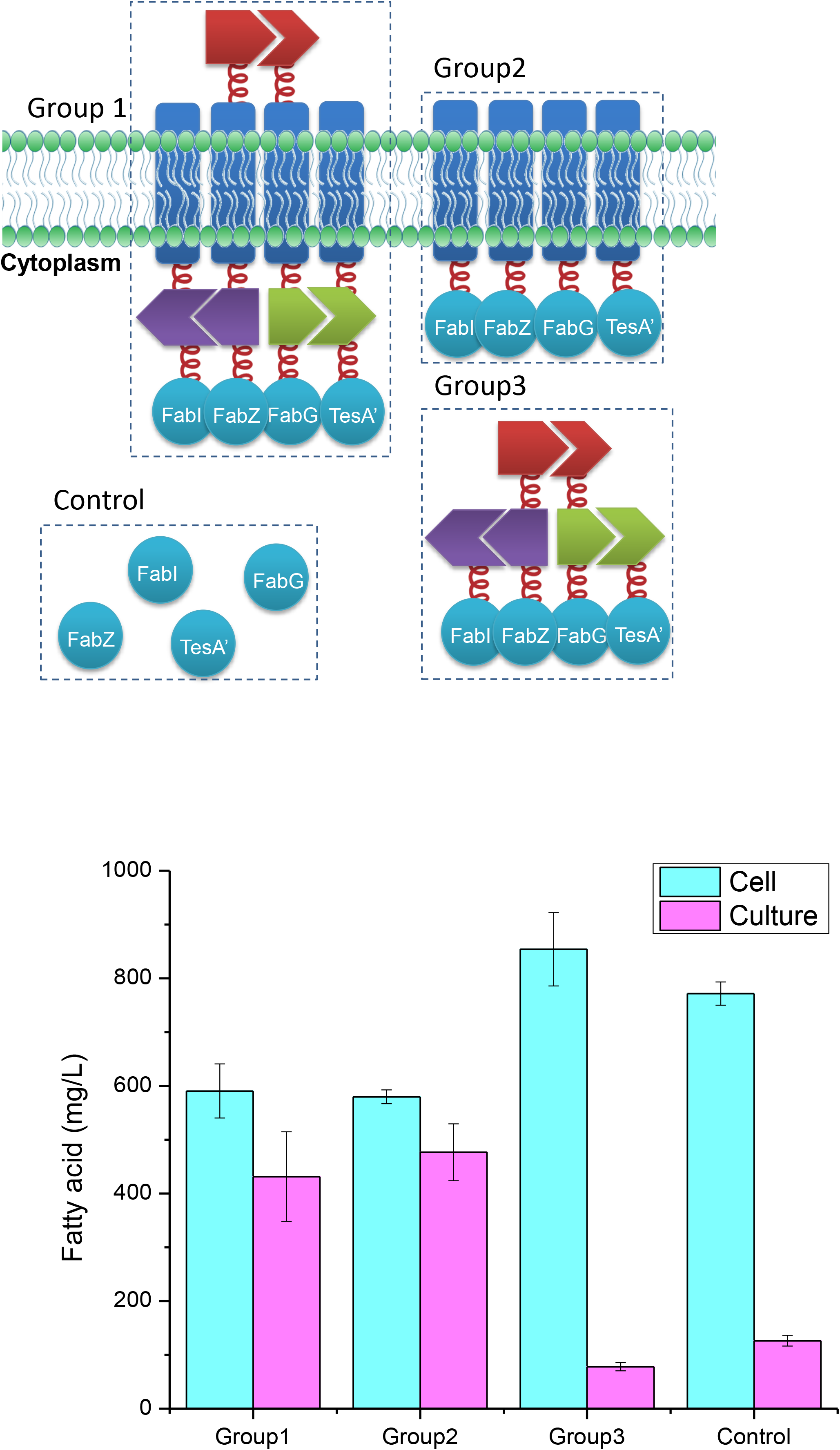
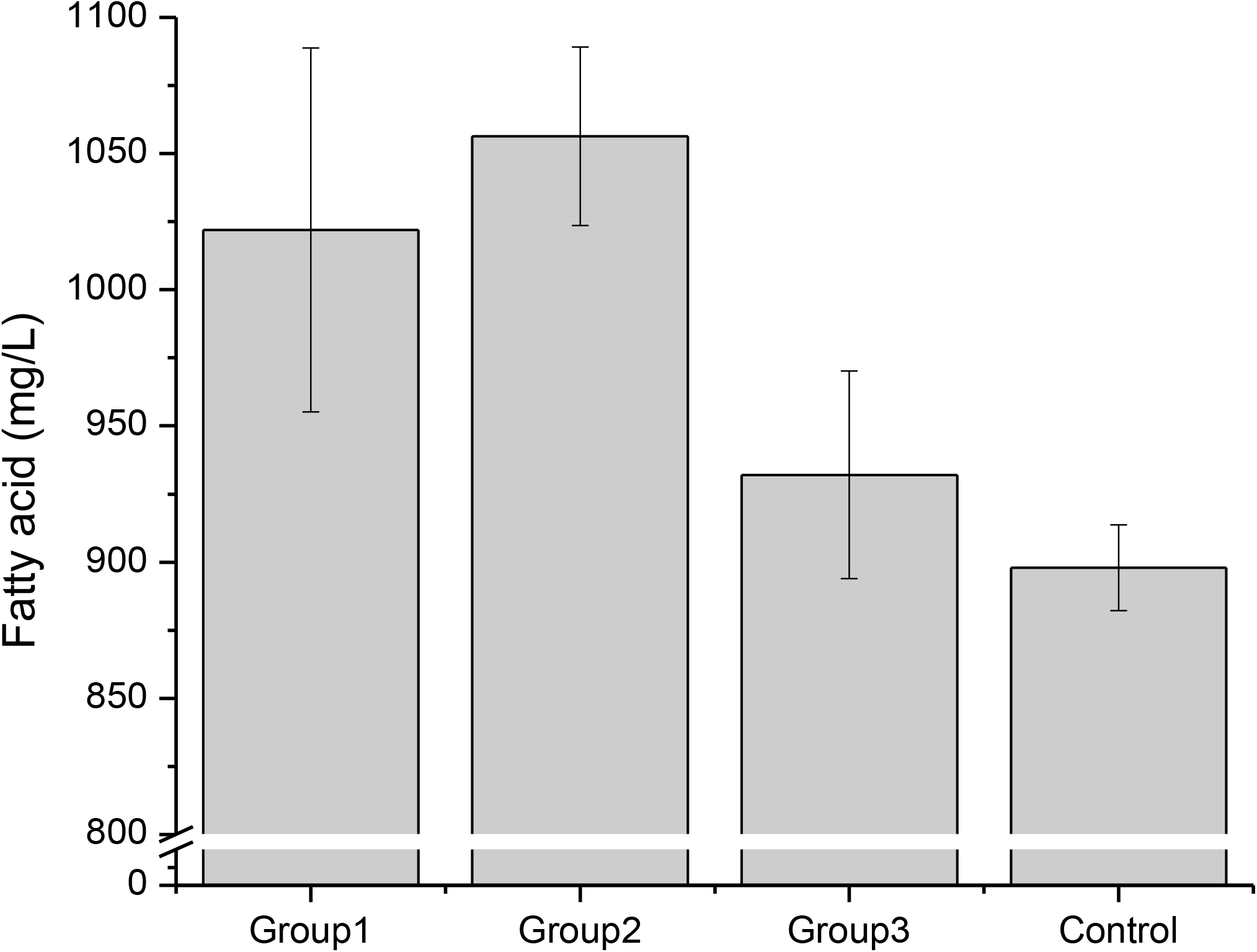
Clustering fatty acid metabolism enzymes on the membrane enhances the products secretion. A) Schematic representation of three strategies to cluster enzymes. Group 1 uses protein interaction domains to cluster target enzymes on the membrane. Group 2 directly localizes target enzymes on the membrane. Group 3 uses protein interaction domains to cluster target enzymes in the cytoplasm. Control group is overexpressed cytoplasm target enzymes. B) Fatty acid yield produced from different groups and from the cell or culture are extracted and compared. C) Total fatty acid from different groups are extracted and compared.

## Discussion

Organisms naturally cluster some related enzymes to improve the efficiency of the whole metabolism pathway and save energy (*1, 2*). Following this idea from nature, many systems have been developed to cluster target enzymes (*5, 6*). The natural scaffolds or artificial scaffolds used in these systems generally need complicated construction process and may have limited slots for enzymes needed to be clustered (*5, 6*). To simplified the clustering system and increased the number of enzymes that can be applied to the system, we noticed that the properties of the cell membrane could meet our goal. First, cell membrane has much smaller space compared to the cytoplasm volume, potentially clustering overexpressed artificial membrane proteins. Second, as a two-dimensional plane, cell membrane has uncountable slots for incoming membrane proteins. Together, we choose to use cell membrane as a scaffold to cluster target enzymes.

To verify this idea, we first designed *E.coli* inner cell membrane anchor construct. Based this construct, we made a control group (Fig. 3A, group 1) to force target enzymes to cluster on the membrane and applied *E.coli* fatty acid synthesis pathway to the system. Comparing to this control group, we found that simply anchoring enzymes on the membrane (Fig. 3B, group 2) showed similar or better yield of the total products. This result indicated that cell membrane can restrict membrane proteins in relative small region compared to the cytoplasm, resulting in the similar effects as clustering proteins. Following this idea, we tried to use fluorescence complementation or fluorescence resonance energy transfer experiments to verify whether simply anchoring on the membrane can cluster proteins. However, we did not observe positive results. Our explanation is that the enzymes anchored on the membrane are close enough to generate clustering effects and enhance the metabolism flux, but are not close enough to be detected by fluorescence complementation or fluorescence resonance energy transfer experiments. Even though, our results show that anchoring enzymes on the membrane can enhance the metabolism flux as described in other artificial scaffold systems (5, 6).

To our surprise, when we analyzed the fatty acid products from cell and from medium, we found that anchoring enzymes on the membrane dramatically increased the products exportation. To verify this phenomenon, we compared four groups of enzymes with different locations (Fig. 4A). The results confirmed that anchoring enzymes on the membrane enhanced the products amount in the medium, indicating more products have been exported. The rate that small molecules randomly diffuse through cell membrane is low and related to the concentration difference between two sides. Our results suggest that when anchoring enzymes on the membrane, the products will accumulate near the cell membrane, resulting in the higher local concentration. This locally increased concentration drives the products molecules diffuse through the cell membrane. Thus, we observed dramatically more products in the medium. Additionally, we noticed that anchoring enzymes on the membrane showed higher yield than clustered enzymes in the cytoplasm. This is possibly because that the products are constantly exported and not accumulated in the cell. As a simple reaction model, accumulated products will inhibit or reverse the reaction. By constantly exporting products, the reaction is driven to the positive direction, leading to the higher total yield.

Together, we show our design of using cell membrane as scaffold to clustering target enzymes to enhance the metabolism flux. The construction process is simplified as fusing target enzymes to the N-terminal or the C-terminal of the membrane anchor protein (Lgt), and the number of the enzymes are not limited (Fig. 5A). Potentially, enzymes can be anchored in the periplasm and utilize substrates from medium to make target products (Fig. 5B). Our design not only shows similar enzymes clustering effects as other artificial scaffolds, but also enhance the products exportation, driving the whole metabolism flux to the positive direction and resulting in further increased final yield compared to the cytoplasm scaffold system.

**Figure 5.**
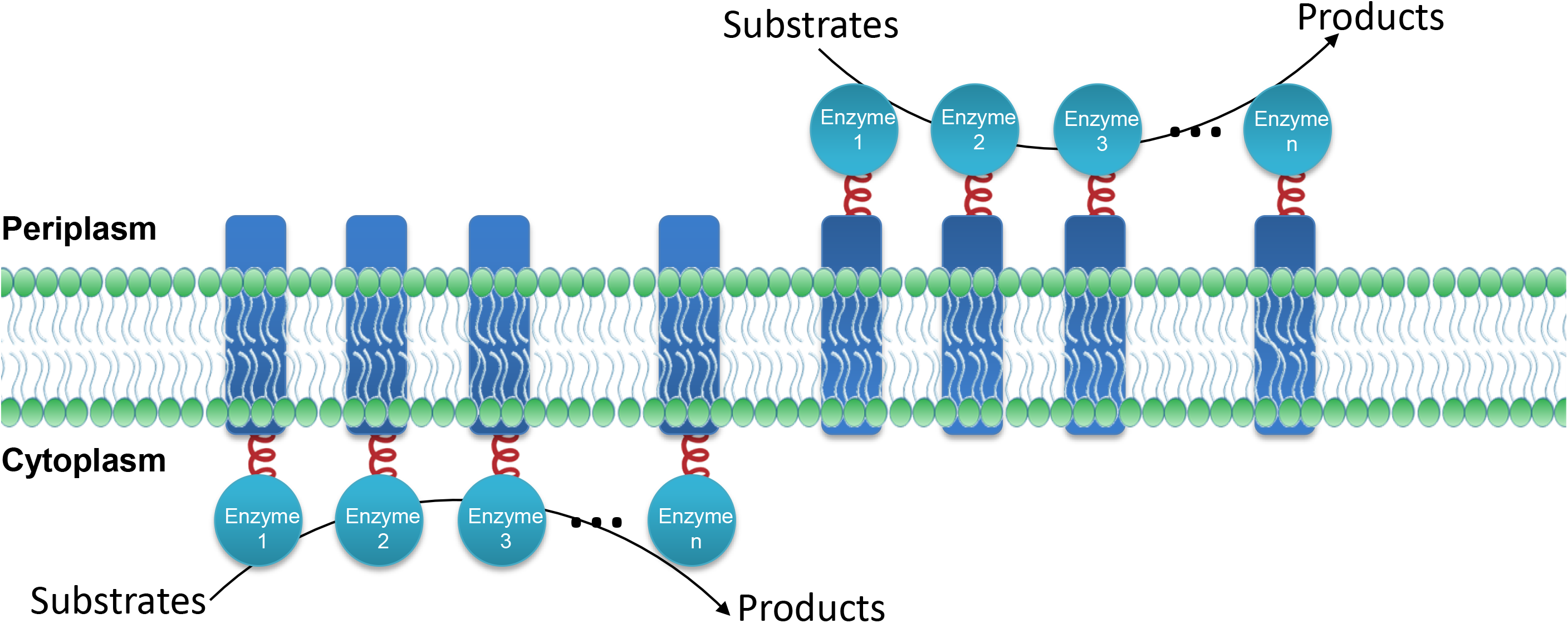
Summary of applications of membrane scaffold system. Potentially, unlimited number of target enzymes can be fused to Lgt and clustered on the membrane. The enzymes can be either presented in the cytoplasm site or the periplasm site. In the cytoplasm site, enzymes can utilize substrates produced in the cell, and the products could pass the cell membrane by diffusion. In the periplasm site, enzymes can utilize the substrates added in the culture and directly release products into the culture.

## AUTHOR CONTRIBUTIONS

Y.W. and G.M. conceived and designed the experiments. Y.W. Y.Q.W., Y.S., H.Q.G., Y.N.Y., R.N.Y., R.X., J.J.W., N.H., Y.H.Z., S.B.Z., and Z.M.J. performed the experiments. Y.W., Y.S.W., and G.M. analyzed the data. Y.S.W., L.H., and G.M. contributed reagents/materials/analysis tools. Y.W. and G.M. wrote the paper.

## ACKNOWLEDGEMENTS

This work was supported by the National Nature Science Foundation of China (31671504, 81421061) and the National Key Technology R&D Program (2012BAI01B09).

